# STAT3 regulates basal cell identity and morphogenesis during early esophageal development

**DOI:** 10.1101/2025.08.01.668154

**Authors:** Secunda W. Kariuki, Yosuke Mitani, Dominique D. Bailey, Gizem Efe, Ved V. Tripathi, Halil Tekin, Kensuke Suzuki, Jianwen Que, Joel Gabre, Ricardo Cruz-Acuña

## Abstract

The transcription factor STAT3 plays broad roles in epithelial biology, yet its function in human esophageal development remains undefined. Using 2D and 3D human induced pluripotent stem cell (hiPSC)-derived platforms, we investigated how STAT3 regulates esophageal epithelial differentiation. We find that STAT3 is dispensable for definitive endoderm and anterior foregut endoderm specification but becomes essential during the transition to esophageal progenitor cells (EPCs). Inhibition of STAT3, via CRISPR-mediated knockout or siRNA, impairs the expression of key EPC and differentiation markers, including *TP63*, and disrupts 3D organoid formation. These defects are accompanied by reduced epithelial proliferation. Notably, STAT3 is highly expressed in human fetal esophageal tissues and hiPSC-derived organoids, while its deletion in the developing mouse esophagus does not affect epithelial architecture, highlighting species-specific differences. Together, these findings identify STAT3 as a critical determinant of basal cell identity and epithelial morphogenesis, revealing a developmental checkpoint in early human esophageal lineage commitment.

## Introduction

The human esophageal epithelium develops from the cranial-most portion of the gut tube called the anterior foregut endoderm (AFE)^1^. By the fourth week of development, differential activation of transcription factors (TFs) along the dorsoventral axis of the AFE dictates whether cells commit to an esophageal or a respiratory fate^2,3^. High Sox2 expression and the absence of Nkx2.1 in dorsal AFE cells promote esophageal lineage commitment and repress respiratory identity ^2,3^. This molecular profile enables dorsal AFE cells to give rise to multipotent fetal esophageal progenitor cells (EPCs), which in turn generate the stratified epithelial cell types that populate the esophagus^4,5^.

By 10 weeks of human gestation, EPCs begin to differentiate into basal cells expressing p63, KRT5, and KRT14, and into suprabasal cells expressing KRT13 and Involucrin^4,5^. This stratification coincides with sustained Sox2 expression during epithelial development. Interestingly, STAT3, a transcription factor with known roles in epithelial differentiation, is highly expressed in both basal and suprabasal layers of the fetal human esophagus (**Fig. 1A**), suggesting a potential regulatory role in esophageal epithelial morphogenesis and lineage specification.

**Figure 1.**
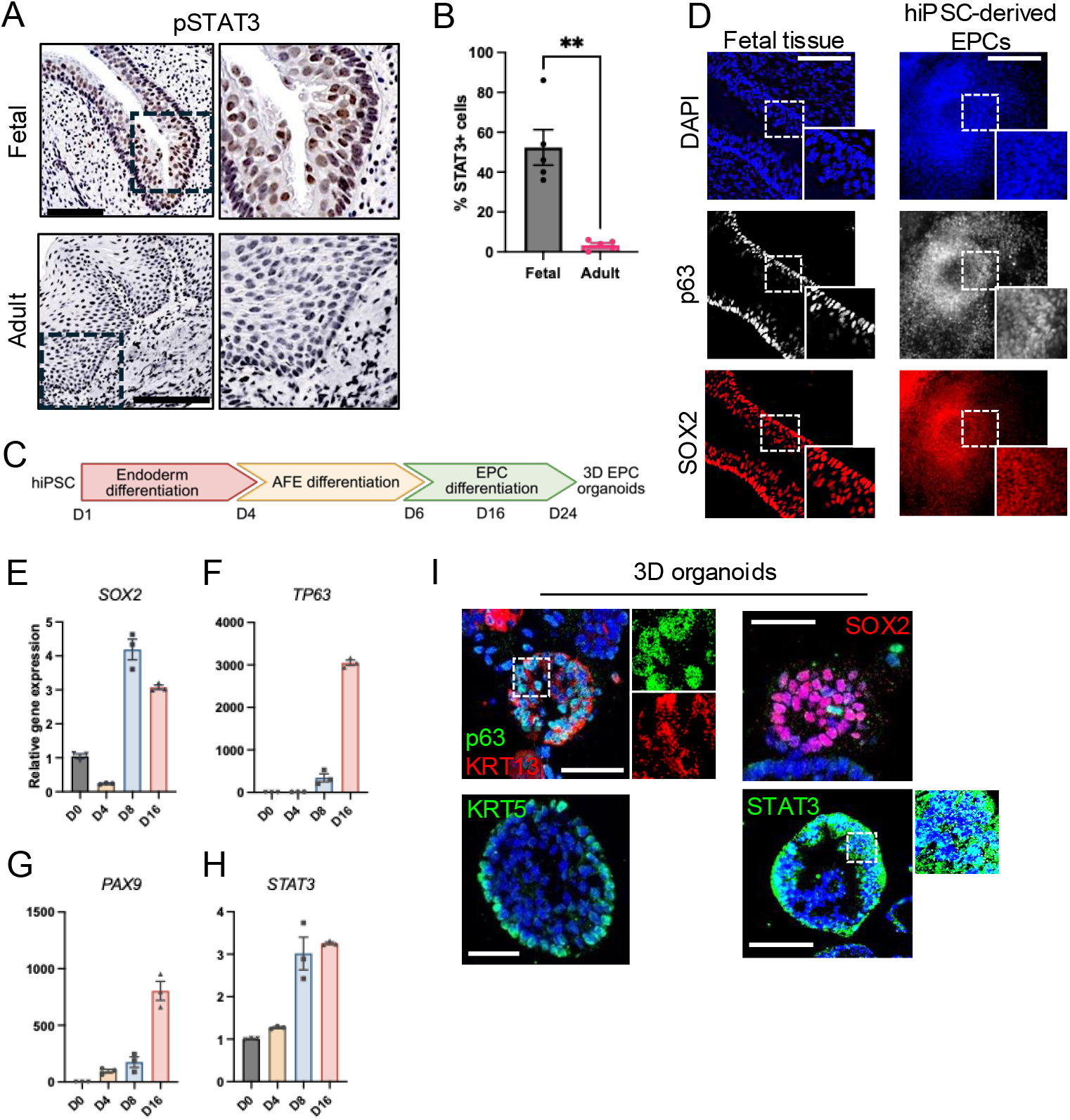
STAT3 is active during human fetal esophageal development and hiPSC-derived esophageal differentiation. (A) Immunohistochemical staining and (B) quantification of phosphorylated STAT3 (pSTAT3) in 10-week human fetal esophageal tissue and adult esophagus. (C) Schematic timeline of directed differentiation protocol used to generate esophageal progenitor cells (EPCs) from BU3 hiPSCs, as previously described^15^. (D) Immunofluorescence staining of fetal esophageal epithelium and hiPSC-derived EPCs for p63, SOX2, and KRT5, with DAPI nuclear counterstain. (E–H) qPCR analysis of (E) *SOX2*, (F) *TP63*, (G) *PAX9*, and (H) *STAT3* mRNA expression during a 16-day differentiation protocol. (I) Immunofluorescence staining of hiPSC-derived 3D esophageal organoids (day 10 post-embedding in Matrigel™) for p63, KRT13, SOX2, KRT5, and STAT3. Scale bars: 100 µm. Graphs: mean ± SEM. (B) n = 6 different biological replicates per condition. Welch’s t-test, two-tailed. **P < 0.01. (E-H) n = 3 wells per condition. All experiments were performed independently at least three times; representative results are shown.

STAT3 is a member of the STAT protein family that mediates transcriptional responses to extracellular cues^6^. Upon activation by Janus kinases (JAKs), phosphorylated STAT3 (pSTAT3) dimerizes, translocates to the nucleus, and regulates genes involved in proliferation, differentiation, and survival^6–8^. Previous studies have shown that STAT3 contributes to gastrointestinal development, including intestinal epithelial cell maturation^9,10^. In rodent embryos, pSTAT3 is highly expressed in the developing lung epithelium^11,12^, further implicating STAT3 in epithelial tissue patterning. However, significant anatomical and histological differences between rodent and human esophagus, such as the presence of submucosal glands and the lack of a keratinized surface in humans^13,14^, limit the translational relevance of rodent studies to human esophageal development. To explore this further, we generated tamoxifen-inducible mouse models with conditional deletion of Stat3 in either p63+ or Krt5+ basal cells. Despite efficient recombination, STAT3 loss did not affect esophageal epithelial architecture by late gestation (**Supplementary Fig. 1**), suggesting species-specific differences in developmental requirements. These findings further justify the use of a human model to dissect the role of STAT3 in esophageal epithelial differentiation.

To address this limitation, we employed a human-specific model based on the directed differentiation of human induced pluripotent stem cells (hiPSCs). This approach enables the generation of relevant esophageal lineages and 3D organoids that recapitulate key features of early epithelial development^15^. Previous studies have used hiPSC-based models to reveal mechanisms of esophageal development, including EPC specification^16^, tracheoesophageal separation^13^, and epithelial morphogenesis^17^. Consistent with these efforts, we differentiated hiPSCs into EPCs that express canonical markers of fetal esophageal development, and formed 3D human esophageal organoids (HEOs) that mimic *in vivo* epithelial architecture. We leveraged this system to investigate how STAT3 regulates EPC lineage commitment, epithelial differentiation, and 3D organization.

Here, we demonstrate that while STAT3 is dispensable for endoderm and AFE differentiation, it is required for EPC commitment, epithelial differentiation, and organoid formation. STAT3 loss impairs expression of EPC-specific markers, reduces proliferation, and limits the ability of cells to generate organized 3D structures. To our knowledge, this is the first study to define the role for STAT3 in early human esophageal epithelial development using hiPSC-derived models.

## Results

### pSTAT3 is enriched during early esophageal development and coincides with epithelial lineage marker expression

STAT3 is known to regulate early epithelial differentiation in multiple tissues, including the lung and intestine^9–12^. To determine whether STAT3 is active during human esophageal development, we examined pSTAT3 protein expression in 10-week-old human fetal esophageal tissue. We observed strong pSTAT3 expression, particularly in the basal layer (**Fig. 1A**). Interestingly, pSTAT3 expression was significantly reduced in the adult esophageal epithelium (**Fig. 1A,B**). These findings suggest a developmental role for STAT3 during epithelial stratification in the fetal esophagus.

To study STAT3 activity *in vitro*, we used a directed differentiation protocol^15^ to derive EPCs from the BU3 human iPSC line **(Fig. 1C)**. Between days 1 and 4, cells were differentiated toward definitive endoderm (CXCR4+, c-KIT+; **Supplemental Fig. 2A**,**B**), followed by BMP and TGF-β inhibition from days 4 to 6 to generate anterior foregut endoderm (AFE)^15,18^. Continued inhibition through day 16 promoted EPC identity, characterized by expression of p63, SOX2, and PAX9 (**Fig. 1D-G**). Cells were then maintained in serum-free differentiation medium until day 24. We observed a significant increase in STAT3 transcription beginning at day 4, which remained elevated through day 16 (**Fig. 1H**), coinciding with the induction of EPC markers. These data suggest that STAT3 expression is temporally associated with EPC differentiation.

**Figure 2.**
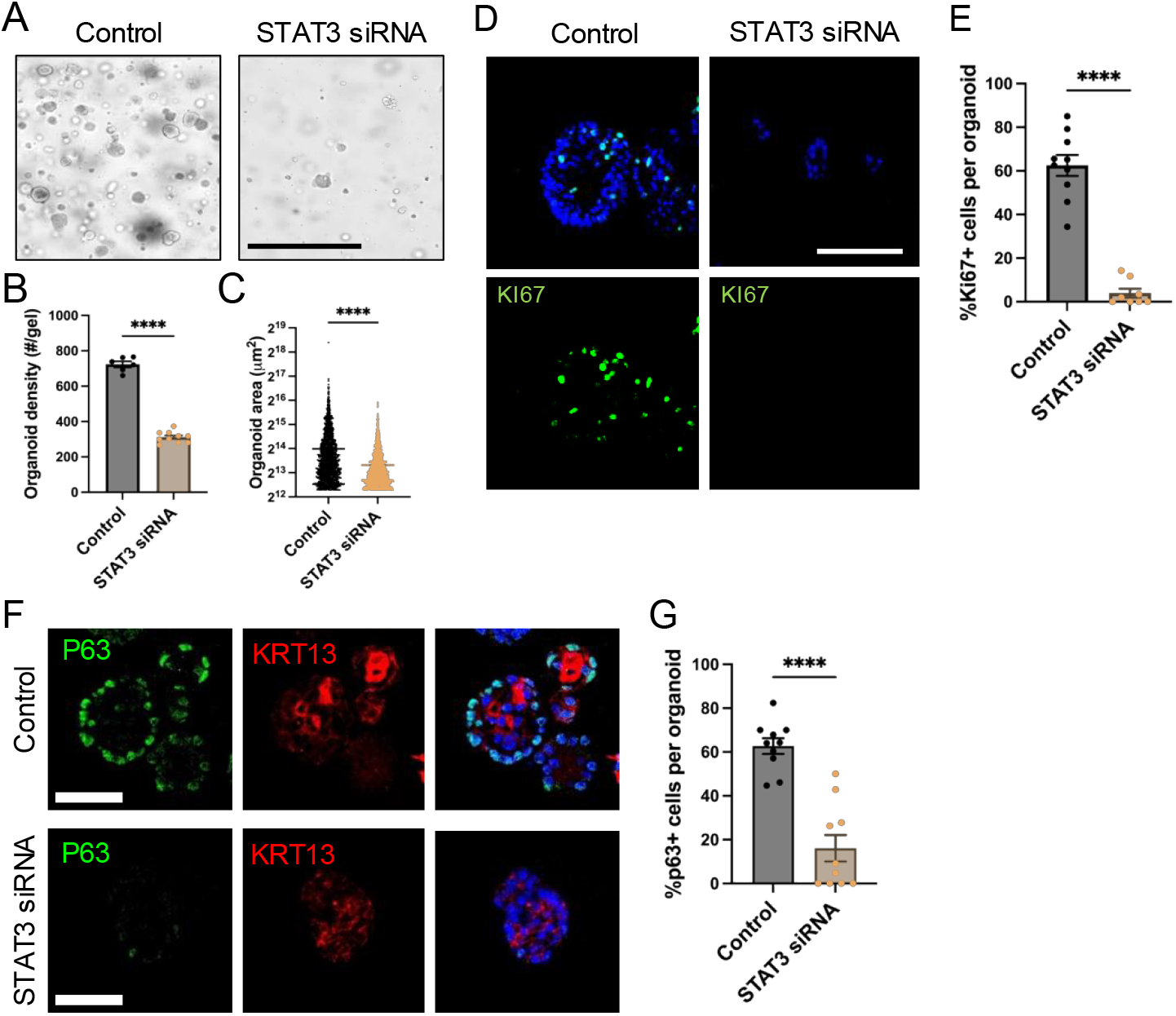
STAT3 regulates organoid formation, proliferation, and epithelial differentiation in esophageal progenitor cells. (A) Representative brightfield images of hiPSC-derived EPC organoids cultured for 10 days in Matrigel™ following transfection with control (scrambled) or STAT3 siRNA. (B, C) Quantification of (B) organoid density (number of organoids per well) and (C) organoid size (area) across conditions. (D) Immunofluorescence staining for Ki67 in EPC-derived organoids at day 10 post-embedding. (E) Quantification of Ki67+ cells per organoid in control vs. STAT3 siRNA conditions. (F) Immunofluorescence staining for p63 and KRT13 in EPC organoids. (G) Quantification of p63+ cells per across conditions. Scale bars: (A) 1 mm, (D,F) 100 µm. Graphs: mean ± SEM. (B) n > 5 wells per condition; (C) n > 500 organoids per condition; (E) n = 10 organoids per condition; (G) n = 10 organoids per condition. Welch’s t-test, two-tailed, ****P < 0.0001 for all comparisons. Experiments were independently performed at least three times; representative results are shown.

To assess STAT3 activity in 3D culture, we embedded day 24 EPCs in Matrigel™ to generate esophageal organoids. After 10 days, these organoids expressed canonical markers of the fetal esophageal epithelium, including P63, KRT13, KRT5, and SOX2, and showed high levels of STAT3 (**Fig. 1I**). These findings indicate that STAT3 is active during esophageal epithelial development in both fetal tissue and hiPSC-derived organoids. Together with prior studies^13^, our data support the fidelity of this organoid system as a platform to model human fetal esophageal development.

### STAT3 inhibition impairs EPC organoid formation, proliferation, and epithelial differentiation

Given the robust expression of STAT3 in fetal esophageal tissue and EPC organoids, we next asked whether STAT3 is functionally required for organoid formation and growth, key indicators of epithelial progenitor potential. To determine this, we transiently inhibited STAT3 expression by transfecting differentiating EPCs with STAT3 siRNA on day 23 of the 24-day protocol. STAT3 knockdown was confirmed by qPCR the following day (**Supplementary Fig. 3A**), and cells were embedded in Matrigel to generate 3D organoids. By day 10 post-embedding, EPCs transfected with STAT3 siRNA formed significantly fewer and smaller organoids compared to controls (**Fig. 2A–C**). Similar results were observed using organoids derived from primary 10-week human fetal esophageal epithelium, in which STAT3 knockdown also reduced 3D organoid size (**Supplementary Fig. 4**). These results indicate that STAT3 activity is necessary for proper organoid growth in both EPCs and primary fetal esophageal cells.

**Figure 3.**
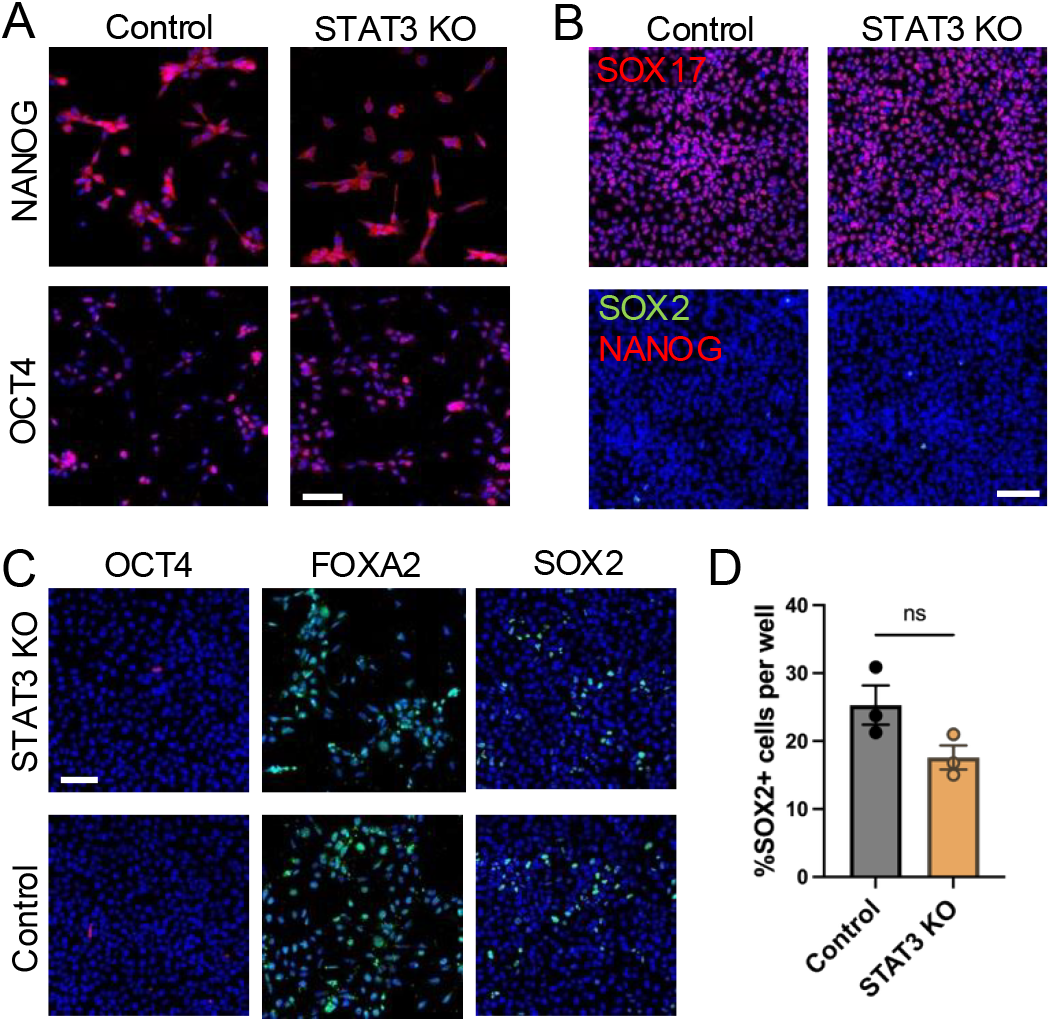
STAT3 is dispensable for definitive endoderm and anterior foregut endoderm differentiation. (A) Immunofluorescence staining of undifferentiated STAT3 knockout (STAT3 KO) and scrambled sgRNA control hiPSCs for OCT4 and NANOG. (B) Immunofluorescence staining of differentiating cells at day 4 for SOX17, SOX2, and NANOG in STAT3 KO and control lines. (C) Immunofluorescence staining at day 6 for anterior foregut marker SOX2 and FOXA2, along with OCT4 to assess loss of pluripotency in STAT3 KO and control lines. (D) Quantification of %SOX2+ cells at day 6 between STAT3 KO and control lines. Scale bars: 100 µm. Graphs: mean ± SEM; n = 3 wells per condition. Welch’s t-test, two-tailed. ns = not significant. All experiments were independently performed three times; representative results are shown.

**Figure 4.**
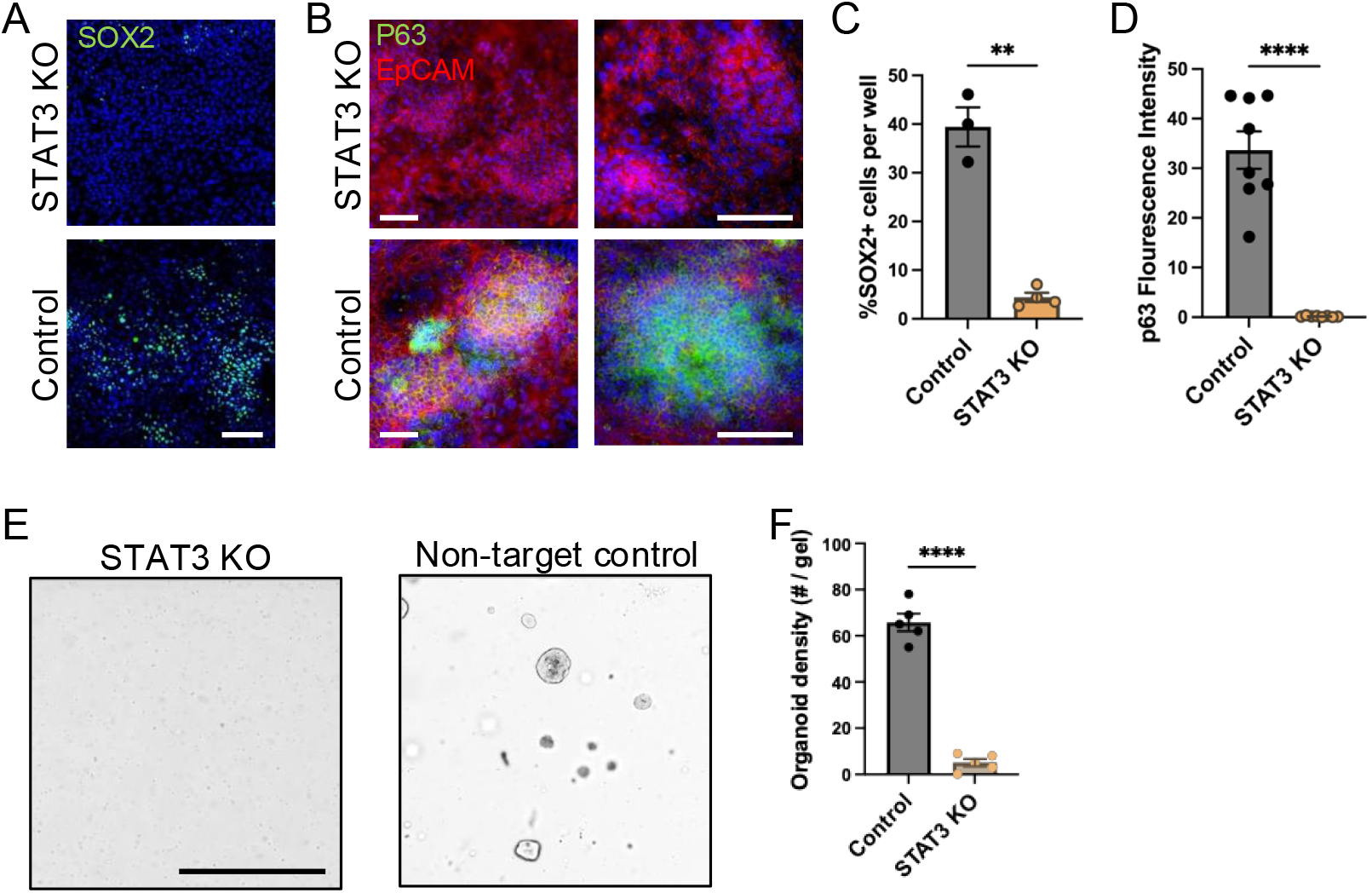
STAT3 is required for esophageal progenitor specification and 3D epithelial organization. (A,B) Immunofluorescence staining at day 16 of differentiation for (A) SOX2, (B) P63 and EpCAM in STAT3 knockout (STAT3 KO) and scrambled sgRNA (non-target) control hiPSC lines. (C,D) Quantification of (C) SOX2+ and (D) P63+ cells per well in STAT3 KO and control lines. (E) Representative brightfield images of 3D esophageal organoids derived from STAT3 KO or control cells, embedded in Matrigel™ on day 24 and cultured for 10 days. (F) Quantification of organoid density (number of organoids per well) in STAT3 KO and control conditions. Scale bars: (A,B) 100 µm, (E) 1 mm. Graphs: mean ± SEM; n > 3 wells per condition. Welch’s t-test, two-tailed. **P < 0.01; ****P < 0.0001. All experiments were performed independently three times; representative results are shown.

To investigate whether reduced organoid growth was due to impaired proliferation or increased apoptosis, we stained organoids for Ki67 on day 10. STAT3 knockdown organoids exhibited significantly fewer Ki67+ cells, as compared to the control group (**Fig. 2D,E**). These data suggest that STAT3 promotes EPC organoid formation by regulating epithelial proliferation.

We next examined whether STAT3 also regulates epithelial differentiation. Compared to control, STAT3 knockdown significantly decreased transcription of basal cell markers (*KRT5, TP63, PAX9*), the suprabasal differentiation marker *KRT13*, and fetal esophageal markers *SOX9* and *KRT7* (**Supplementary Fig. 3B-G**). Protein expression of P63 and KRT13 was also markedly reduced in STAT3 siRNA organoids (**Fig. 2F,G**). Together, these findings highlight STAT3 as a critical regulator of early esophageal epithelial development, linking progenitor identity to proliferative capacity and morphogenetic competence in 3D culture.

### STAT3 is not required for endoderm and anterior foregut endoderm linage commitment

Because STAT3 is activated early in development and regulates differentiation in other embryonic tissues, we first asked whether it is required for early germ layer specification. To test this, we generated a STAT3 knockout (STAT3 KO) hiPSC line using CRISPR/Cas9 targeting of STAT3 in the BU3 background. A control line was established using a scrambled (non-target) sgRNA. Successful knockout was confirmed by western blot (**Supplementary Fig. 5**), and both lines maintained expression of the pluripotency markers OCT4 and NANOG (**Fig. 3A**), indicating that loss of STAT3 does not impair stem cell identity^19^.

We next assessed whether STAT3 KO cells could differentiate into definitive endoderm. From day 1 to 4 of the differentiation protocol, both STAT3 KO and control lines showed robust induction of the definitive endoderm marker SOX17, along with downregulation of the pluripotency markers SOX2 and NANOG (**Fig. 3B**)^15,20^. Flow cytometry at day 4 further revealed high and comparable surface expression of CXCR4 and c-KIT in both lines, confirming successful generation of definitive endoderm (**Supplementary Fig. 2C-F**)^18^.

To determine whether AFE specification was also preserved, we continued differentiation to day 6 and examined expression of the AFE markers SOX2 and FOXA2^1^. As expected, SOX2 was re-expressed in both STAT3 KO and control lines at this stage, consistent with its known role in anterior foregut patterning^1^. Both STAT3 KO and control lines exhibited strong expression of these markers (**Fig. 3C–D**), and residual OCT4 signal was negligible, confirming efficient lineage progression. These findings indicate that STAT3 is dispensable for differentiation into definitive endoderm and AFE. To our knowledge, this is the first demonstration that STAT3 is not required for early germ layer specification in human pluripotent stem cells, highlighting that STAT3 function is required after endoderm and AFE specification, during esophageal epithelial lineage commitment.

### STAT3 regulates esophageal progenitor cell specification, differentiation, and 3D organization

Having established that STAT3 is not required for endoderm or AFE specification, we next asked whether it plays a role in the subsequent transition to esophageal progenitor cells (EPCs), a critical step in stratified epithelial development. To test this, we continued differentiation of STAT3 KO and control hiPSC lines from AFE to EPCs (days 6–16). By day 16, STAT3 KO cells exhibited significantly reduced expression of the EPC markers P63 and SOX2 compared to controls, while expression of EpCAM remained high in both lines, indicating that epithelial identity was retained (**Fig. 4A–D**). Cell density and morphology remained similar between groups, suggesting that STAT3 loss specifically impaired progenitor specification rather than overall viability or epithelialization. EPC differentiation was repeated using three validated STAT3 KO clones with consistent results (data not shown).

To further investigate the transcriptional impact of STAT3 loss during EPC specification, we performed bulk RNA sequencing of STAT3 KO and control cells at day 16 of differentiation. While TP63 expression was modestly reduced, interestingly, other canonical EPC markers such as PAX9 and KRT5 were not significantly altered. However, gene ontology (GO) analysis of downregulated transcripts revealed strong enrichment for pathways associated with epithelial identity and stratification (**Supplementary Fig. 6A-C**). Upregulated genes were enriched for neuronal and synaptic processes, which we interpret as evidence of lineage instability and transcriptional derepression. This interpretation is consistent with previous reports that failure to maintain p63+ EPCs in hPSC-derived cultures results in overgrowth of neuroectodermal cells^15^. Notably, several of the most upregulated genes in STAT3 KO EPCs, including WNT8B^21^, FAM107A^22^, and TRIM58^23^, are associated with developmental signaling and neuronal cytoskeletal remodeling (**Supplementary Fig. 6D**), further supporting this model.

To further validate the requirement for STAT3 in EPC specification, we performed an independent experiment using STAT3 siRNA. AFE cells (day 6) from the parental hiPSC line were transfected with STAT3 siRNA, and gene expression was analyzed on days 16 and 24 (**Supplementary Fig. 7A**). STAT3 levels remained significantly suppressed in the siRNA group across both timepoints (**Supplementary Fig. 7B**). In contrast to the RNA sequencing data, expression of EPC commitment markers (*SOX2, PAX9, TP63*) and differentiation markers (*KRT5, KRT13*) was significantly reduced following STAT3 knockdown (**Supplementary Fig. 7C–G**). These results reinforce that STAT3 is necessary for the transition from AFE to EPCs and for expression of early esophageal epithelial identity.

Next, to further validate whether STAT3 is also required for 3D epithelial organization, we embedded day 24 cells from STAT3 KO and control lines in Matrigel™ and analyzed 3D organoid formation after 10 days. As with siRNA knockdown (**Fig. 2**), STAT3 KO cells formed significantly smaller and fewer organoids compared to controls (**Fig. 4E–F**), confirming that STAT3 is required for proper 3D organization and self-assembly of esophageal progenitors. Together, these findings demonstrate that STAT3 is essential for establishing esophageal progenitor identity and enabling their self-organization into 3D epithelial structures, highlighting a specific developmental checkpoint where STAT3 becomes functionally required.

## Discussion

In this study, we define a novel role for STAT3 in regulating the early differentiation and organization of human esophageal epithelium. Using a human iPSC-derived model, we demonstrate that STAT3 is not required for definitive endoderm or anterior foregut endoderm specification, but becomes essential during the transition to esophageal progenitor cells (EPCs). Loss of STAT3 impairs EPC differentiation, suppresses expression of key epithelial markers including TP63, and disrupts 3D organoid formation, indicating that STAT3 orchestrates both cell identity and morphogenetic potential during early esophageal development.

Our findings position STAT3 as a developmental regulator acting downstream of AFE, where it appears to coordinate epithelial fate acquisition and self-organization. This temporally specific requirement contrasts with its established roles in other endoderm-derived organs, such as the intestine, where STAT3 promotes epithelial maturation and barrier function^9,10^. In the lung, STAT3 activity has been shown to regulate epithelial integrity and differentiation during embryogenesis^11,12^. That STAT3 is dispensable during early lineage specification in our model, but required for subsequent epithelial differentiation, suggests organ-and stage-specific deployment of its transcriptional targets.

A central finding of this work is the significant reduction in TP63 expression following STAT3 inhibition. p63 is a master regulator of basal epithelial cell fate and is essential for the formation and maintenance of stratified squamous epithelia^24,25^. In both STAT3 knockout and siRNA conditions, we observed reduced p63 at the transcriptional and protein level, alongside downregulation of other basal (e.g., KRT5, PAX9) and suprabasal (e.g., KRT13) markers. These findings are consistent with work in the skin, where STAT3 promotes keratinocyte proliferation and supports basal cell renewal^26,27^. Prior studies have shown that p63 and SOX2 cooperatively regulate gene expression through AP-1-associated elements^28^, and that SOX2 and STAT3 cooperate in the malignant transformation of foregut epithelial cells^29^. Together, these data raise the possibility that STAT3, SOX2, and p63 function as a regulatory network during esophageal epithelial development and disease.

Transcriptomic profiling of STAT3 KO EPCs provided further insight into how STAT3 regulates epithelial identity at the systems level. GO enrichment analysis of downregulated transcripts revealed loss of epithelial adhesion and differentiation pathways, whereas upregulated transcripts were enriched for neuronal and synaptic gene programs. This expression shift likely reflects transcriptional derepression rather than fate switching, consistent with prior work showing that neuroectodermal cells emerge in hPSC-derived EPC cultures when epithelial identity is not maintained^15^. Of note, while EpCAM expression was retained in STAT3 KO EPCs, this marker broadly labels epithelial-like states and does not distinguish cells with stable foregut identity from those undergoing transcriptional drift. These results, together with our siRNA-based validation, support the role of STAT3 in stabilizing epithelial lineage commitment and repressing alternative transcriptional states during esophageal development.

In addition to its role in transcriptional regulation, STAT3 also appears to support epithelial proliferation and morphogenesis. The phenotypic consequences of STAT3 loss—in particular, the reduction in 3D organoid size and formation—suggest that STAT3 not only supports molecular identity but is also required for epithelial proliferation. We found decreased Ki67 expression in STAT3-deficient organoids, implicating a proliferation-specific role. This aligns with prior studies linking STAT3 to cell cycle progression and stem cell expansion in other epithelial systems^30,31^. Notably, our results in both hiPSC-derived EPCs and fetal esophageal organoids emphasize that this role is not model-specific, but instead reflects a conserved requirement for STAT3 activity in esophageal epithelial growth.

Interestingly, our *in vivo* analysis using tamoxifen-inducible mouse models revealed that conditional deletion of Stat3 in Krt5+ or p63+ epithelial progenitors did not disrupt esophageal epithelial architecture at E15.5. These findings underscore a key difference between murine and human esophageal development, and support prior reports of species-specific epithelial responses to STAT3 signaling^5,13,14^. Rodent esophagus is highly keratinized and lacks submucosal glands, features that differ markedly from the human tissue and may account for the lack of phenotype observed in Stat3 conditional knockout embryos. This reinforces the need for human-specific developmental models, such as the one used here, to elucidate molecular mechanisms underlying esophageal morphogenesis.

Together, our findings reveal that STAT3 is required for early esophageal epithelial differentiation by regulating basal identity, proliferation, and 3D organization. By positioning STAT3 upstream of p63 and epithelial stratification, this study identifies a critical developmental checkpoint that may also have relevance for disease contexts in which basal cell plasticity or epithelial regeneration is disrupted. Future studies should investigate the direct transcriptional targets of STAT3 in EPCs and define how STAT3 may integrate with other lineage-specifying pathways, such as Notch and Wnt, during esophageal development.

## Materials and Methods

### Mouse models and embryo analysis

All animal procedures were approved by the Columbia University Institutional Animal Care and Use Committee (IACUC protocol: AC-AABI2550). Krt5-CreER; Stat3^fl/fl^; Rosa26^LSL-TdTomato^ and p63-CreER; Stat3^fl/fl^; Rosa26^LSL-TdTomato^ mice were maintained on a C57BL/6J background. Pregnant females were administered tamoxifen (0.2 mg/g body weight in sunflower oil, intraperitoneally) on embryonic days E9.5, E11.5, and E13.5. Embryos were harvested at E15.5. Recombination was confirmed by TdTomato (**Supplementary Tables 1 and 2**) fluorescence in the esophageal epithelium. Paraffin-embedded tissue sections were processed for hematoxylin and eosin (H&E) staining or immunofluorescence using antibodies against p63 and KRT5. Epithelial thickness and luminal area were quantified from transverse sections using ImageJ (NIH). Five embryos were analyzed per group.

### hiPSC culture

The BU3 NGST hiPSC line, which contains a knock-in NKX2.1-GFP reporter, was obtained from the Kotton Laboratory (Center for Regenerative Medicine, Boston University). Cells were cultured in mTeSR Plus medium (STEMCELL Technologies) on Matrigel-coated plates (Corning, 354234) under hypoxic conditions (37°C, 5% CO_2_, 5% O_2_). For passaging, cells were dissociated using StemMACS Passaging Solution XF (Miltenyi Biotec, 130-104-688) and mechanically fragmented into small colonies. Cells were split at a 1:6 to 1:20 ratio and passaged every 5–7 days.

### CRISPR/Cas9-mediated STAT3 knockout in hiPSCs

To generate STAT3 knockout (STAT3 KO) hiPSCs, the BU3 NGST line was nucleofected with a ribonucleoprotein (RNP) complex containing 2.5 µg of SpyFi Cas9 protein (Aldevron, 9214) and two synthetic gRNAs (Synthego): GAGAUUAUGAAACACCAAAG and CAUUCGACUCUUGCAGGAAG. Nucleofection was performed using the Amaxa 4D Nucleofector X-Unit (Lonza, AAF-1003X) and the P3 Primary Cell Kit (Lonza, V4XP-3024), with program CB-150. Forty-eight hours post-nucleofection, individual colonies from the mixed population were manually picked into 96-well plates and cultured in mTeSR Plus medium supplemented with 10% CloneR (STEMCELL Technologies). Colonies were expanded for 2 weeks, with medium changes every other day. Genomic DNA was extracted to screen for STAT3-targeted indels. Clones with confirmed homozygous indels were validated by qRT-PCR and Western blot for complete loss of STAT3 expression. EPC differentiation was repeated using three validated STAT3 KO clones, all of which showed consistent phenotypes; representative data are shown. Primer sequences are listed in **Supplementary Table 3**.

### Differentiation and sorting of hiPSC-derived esophageal cells

Differentiations were adapted from previously described protocols^15^. hiPSCs were seeded at 1 × 10^6^ cells/well in 6-well plates (Thermo Scientific, 14-832-11) and cultured for 24 hours in mTeSR media with 10 µM Y-27632 (Selleckchem, S1049). From days 1– 3, cells were treated with the STEMdiff Definitive Endoderm Kit (STEMCELL Technologies, 05110) to induce endoderm. Day 4 definitive endoderm identity was confirmed by FACS for CXCR4 and c-KIT. Cells were passaged using Gentle Cell Dissociation Reagent (STEMCELL Technologies, 100-0485) at a 1:2–1:3 ratio. From days 4–6, cells were cultured in serum-free differentiation (SFD) medium supplemented with 10 µM SB431542 (Selleckchem, S1067), 200 ng/mL Noggin (R&D Systems, 6057-NG-01M/CF), and 100 ng/mL hEGF (Peprotech, AF-100-15-100). On day 7, cells were transferred to normoxic conditions (21% O_2_). From days 7–16, hEGF concentration was increased to 200 ng/mL in SFD medium. From days 16–24, SFD medium was supplemented only with hEGF at 200 ng/mL. SFD medium was prepared using IMDM (375 mL, Corning, 10-016-CV) and Ham’s F-12 (125 mL, Corning, 10-080 CV), supplemented with B27 (1×, Thermo Fisher, 17504044), N2 (1×, Thermo Fisher, 17502048), Glutamax (1×, Thermo Fisher, 35050061), 7.5% BSA (.05%, Thermo Fisher, 15260037), Primocin (0.2 µg/mL, InvivoGen, ant-pm-1), L-Ascorbic Acid (50 µg/mL, Sigma-Aldrich, A4544), and Monothioglycerol (0.5 mM, Sigma-Aldrich, M6145). On days 16 or 24, cells were dissociated with Accutase (Fisher Scientific, NC1793126) and FACS-sorted for EpCAM+, NKX2.1-GFP– cells using BD FACSAria III. Lists of all antibodies used are included in **Supplementary Tables 1 and 2**.

### 3D culture of hiPSC-derived esophageal organoids

To establish 3D organoid culture, sorted EpCAM+ cells (25,000) were embedded in 50uL of Matrigel in 24-well plates (Corning, 3526) and cultured for 10-14 days at normoxic conditions (21% O_2_; 5% CO_2_). Organoid culture medium consisted of SFD medium supplemented with 10 µM Y-27632, 100 ng/mL Noggin, 10 µM SB431542, 3 µM CHIR99021 (Tocris, 4423), 20 ng/mL FGF2 (R&D Systems, 233-FB), and 200 ng/mL hEGF.

### Fetal esophageal organoid generation and culture

Human fetal esophageal tissue was obtained from the Laboratory of Developmental Biology at the University of Washington under IRB protocol AAAU0544. To establish fetal esophageal organoid lines, epithelial cells from 10-week human fetal esophagus were digested using Dispase (Corning, 354235) and 0.25% Trypsin-EDTA (Gibco, 25200056). Enzymes were neutralized with soybean trypsin inhibitor (Sigma, T9128), and cells were filtered through a 100 µm strainer and plated at 5,000–10,000 cells per 50 µL Matrigel. Organoids were expanded in 50% L-WRN conditioned medium and 50% Advanced DMEM/F12 (Thermo), supplemented with GlutaMAX, HEPES, N2, B27, NAC, CHIR99021, EGF, A83-01, SB202190, Gastrin, Nicotinamide, Y-27632, Gentamicin, and antibiotic-antimycotic. For passaging, domes were disrupted in cold PBS, digested with TrypLE Express (Fisher Scientific, 12-605-010), neutralized, and replated. Organoids were passaged every 7 to 10 days and cultured at normoxic conditions (21% O_2_; 5% CO_2_).

### STAT3 inhibition

STAT3 siRNA (Santa Cruz, sc-29493) or control siRNA (MilliporeSigma, SIC003) was delivered at 10 µM using Lipofectamine RNAiMAX (Invitrogen, 13778150), according to manufacturer’s instructions. For chemical inhibition, 10 µM Stattic (Selleckchem, S7024) was added to organoid medium for 8 days, with media changes every 2–3 days.

### Western Blot

Cells were lysed at ∼80% confluency using lysis buffer (20 mM Tris-HCl, 150 mM NaCl, 1 mM EDTA, 1 mM EGTA, 1% Triton X-100, 2.5 mM sodium pyrophosphate, 1 mM β-glycerophosphate, 1 mM Na_3_VO_4_, and 1× protease inhibitor cocktail; Roche). Protein concentrations were measured using the Bradford Assay Dye Reagent Concentrate (Bio-Rad, 5000006). Proteins (25 µg per sample) were resolved using 4–15% Mini-PROTEAN TGX Precast Gels (Bio-Rad, 4561083) and transferred to nitrocellulose membranes (Bio-Rad, 1704159). Membranes were blocked with Odyssey Blocking Buffer (LICOR, 927-40000), incubated with primary antibodies overnight at 4°C, and secondary antibodies for 2 hours at room temperature. Detection was performed using the Bio-Rad ChemiDoc system. Lists of all antibodies used are included in **Supplementary Tables 1 and 2**.

### Immunofluorescence and immunohistochemistry staining

Fixed paraffin sections underwent antigen retrieval in 10 mM citrate buffer (pH 6.0) and were blocked in 5% donkey serum. Samples were incubated with primary antibodies overnight at 4°C and secondary antibodies (1:1000) for 30 minutes at 37°C. DAPI (Vector Laboratories, H-1500) or Hoechst 33342 (Thermo Fisher, H3570) was used as a nuclear stain. For IHC, endogenous peroxidases were quenched using 3% H_2_O_2_ (Sigma-Aldrich H1009), followed by Avidin (Sigma-Aldrich A9275), Biotin (Sigma-Aldrich B4501), and StartingBlock Blocking Buffer (Thermo Fisher 37539). Sections were incubated with primary antibodies, biotinylated secondaries (Vector BA-2001), ABC reagent (Vector PK-6100), and developed using DAB substrate (Vector SK-4100). For immunocytochemistry staining of differentiating hiPSCs, cells were cultured in µ-Plate 24 Well Black ID 14mm ibiTreat plates (ibidi, 82421) and fixed in the plates with 4% (w/v) paraformaldehyde at room temperature for 15-30 minutes. Cells were blocked with 5% donkey serum and incubated with primary antibodies overnight at 4°C. Secondary incubation was performed for 2 hours at room temperature. Hoechst 33342 was used as a nuclear stain. Lists of all antibodies used are included in **Supplementary Tables 1 and 2**.

### Image acquisition and quantification

Immunofluorescence and H&E images were acquired using a Keyence BZ-X800 microscope. Quantification of positive signal was completed using the Keyence analytical algorithms and ImageJ software. Brightfield images of organoids were acquired using the Celigo Image Cytometer. Quantification of organoid size (area) and density was performed using the Celigo’s analytical algorithms.

### RT-qPCR

RNA was isolated using the RNAqueous Total RNA Isolation Kit (Invitrogen, AM1912), and cDNA was synthesized with the High-Capacity cDNA Reverse Transcription Kit (Thermo Fisher, 43-688-13), according to the manufacturer’s instructions. qPCR was performed on an Applied Biosystems 7500 Real Time PCR System using SYBR Green PCR Master Mix (Applied Biosystems). Primer sequences are listed in **Supplementary Table 3**. All data were normalized to YWHAZ using the ΔΔCt method.

### RNA sequencing and analysis

Total RNA was isolated using the RNAqueous™ Phenol-Free Total RNA Isolation Kit (Invitrogen, AM1912) following the manufacturer’s instructions. RNA samples were submitted to Azenta Life Sciences (USA) for bulk RNA sequencing. Raw sequencing data were processed using Azenta’s standard bioinformatics pipeline, which includes quality control, alignment to the reference genome, transcript assembly, and differential expression analysis. Differentially expressed genes were identified using the DESeq2 package. Genes with an adjusted p-value (Padj) < 0.05 and |log2 fold change| > 1 were considered significantly differentially expressed. Differentially expressed gene sets were used for gene ontology (GO) enrichment analysis using the PANTHER Overrepresentation Test (http://geneontology.org/), and representative GO terms were visualized as bar plots in GraphPad Prism 6.0.

### Statistics and reproducibility

All experiments were performed independently at least three times under similar conditions, except experiments shown in Supplemental Figures 3 and 6, which were performed twice, and Supplementary Figure 4, which was performed once. Data shown are of 1 experiment representative of all independent experiments performed under similar conditions. A biological replicate was defined as an independent differentiation or experimental setup performed on a different day. A technical replicate was defined as a separate well or Matrigel dome within the same experiment. EPC differentiation was performed using three independently derived STAT3 KO hiPSC clones, all of which showed consistent phenotypes; representative data are shown. Data analysis was performed in a blinded fashion where applicable. All immunofluorescence or immunohistochemistry images shown are representative of at least 20 images that were stained and imaged for each specific marker per experimental group for each independent experiment. All statistical analyses were performed using GraphPad Prism 6.0. For statistical comparisons between 2 groups, Welch’s *t* test (or Mann-Whitney *U* test for nonparametric data) was used, and among more than 2 groups, 1-way ANOVA (or Kruskal-Wallis test for nonparametric data) was used. P < 0.05 was considered statistically significant.

## Data availability

All data supporting the findings of this study are available from the corresponding author upon reasonable request.

## Author Contributions

Conceptualization: R.C.-A., J.T.G., J.Q.

Methodology: S.W.K., G.E., V.V.T., Y.M., H.T., D.D.B.

Investigation: S.W.K., G.E., V.V.T., Y.M., H.T., D.D.B.

Validation: S.W.K.

Formal Analysis: S.W.K., R.C.-A.

Visualization: S.W.K., R.C.-A.

Writing – Original Draft: S.W.K., R.C.-A.

Writing – Review & Editing: All authors Supervision: R.C.-A.

Corresponding Author: R.C.-A. designed and supervised the study and oversaw project execution and manuscript preparation.

## Declaration of Interests

The authors declare no competing interests.

## Acknowledgements

This research was supported by the National Institutes of Health (NIH), including the National Institute of Diabetes and Digestive and Kidney Diseases (NIDDK) K01DK133620 (to R.C.-A.), K08DK128603 (to D.D.B.), and the Harold Amos Medical Faculty Development Award from the Robert Wood Johnson Foundation (to D.D.B.); the National Cancer Institute (NCI) P01CA098101 (to R.C.-A. and J.T.G.), P01CA098101-20S1 (to S.W.K.), and R01CA272903 (to J.T.G.); and the Columbia University Digestive and Liver Diseases Research Center (CU-DLDRC) grant P30DK132710.

We acknowledge the Herbert Irving Comprehensive Cancer Center’s shared resources, including the Molecular Pathology, Flow Cytometry, Confocal and Specialized Microscopy, Organoid and Cell Culture Core, and Stem Cell Core Facilities at Columbia University Irving Medical Center. Human fetal tissue was provided by the Laboratory of Developmental Biology at the University of Washington under IRB protocol AAAU0544, supported by NIH Award Number 5R24HD000836 from the Eunice Kennedy Shriver National Institute of Child Health and Human Development.

We thank Dr. James M. Wells for his early conceptual input.

## Notes

### Competing Interest Statement

The authors have declared no competing interest.

