## Supplementary Data for "STAT3 regulates basal cell identity and morphogenesis during early esophageal development"

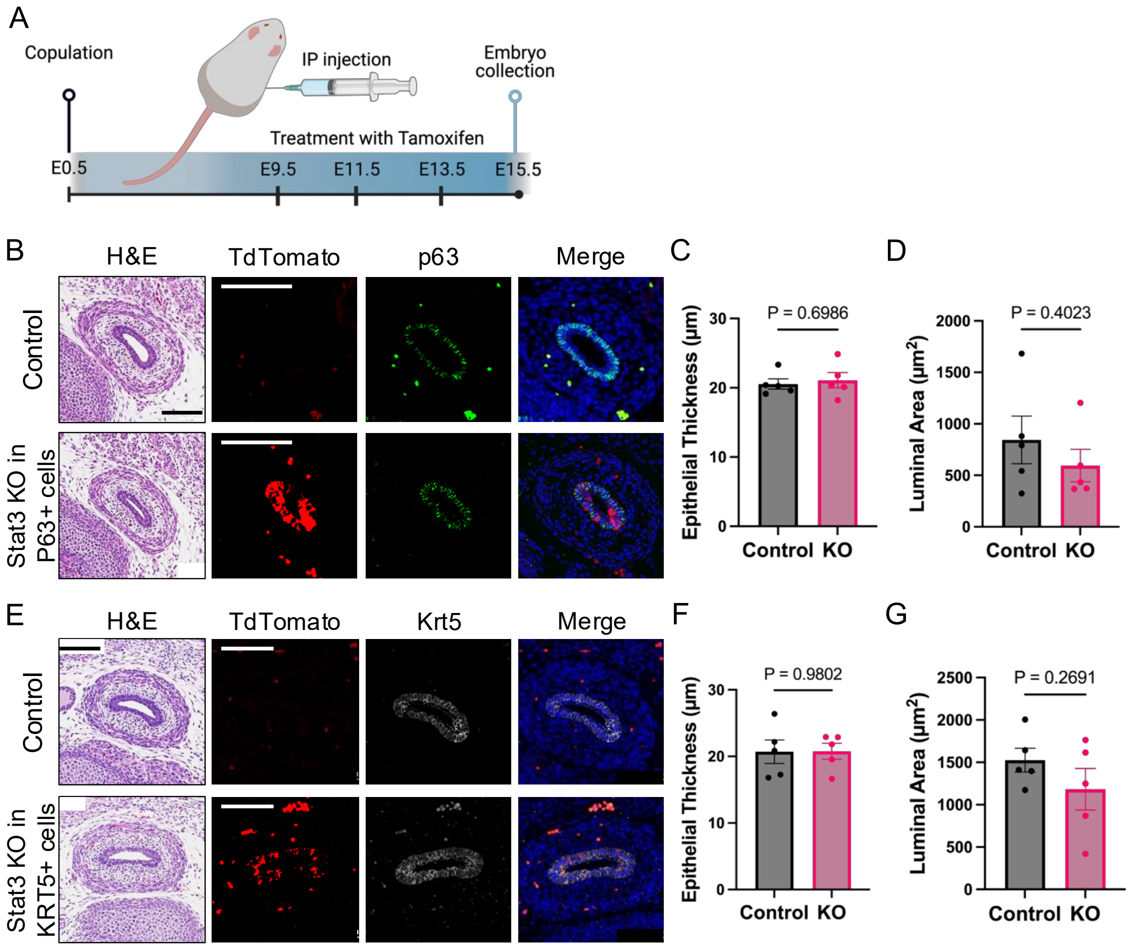


**Supplementary Figure 1. Conditional deletion of Stat3 in embryonic mouse esophagus does not disrupt epithelial development.** (A) Schematic of the tamoxifen-inducible conditional knockout strategy used to generate Krt5-CreER; Stat3$^{fl/fl}$; Rosa26$^{LSL\text{-}TdTomato}$ and P63-CreER; Stat3$^{fl/fl}$; Rosa26$^{LSL\text{-}TdTomato}$ embryos. Pregnant mice received intraperitoneal tamoxifen at three timepoints, and embryos were harvested at E18.5. (B) Representative transverse sections of P63-CreER; Stat3$^{fl/fl}$ (Stat3 KO) embryos stained with H&E and immunofluorescence for TdTomato (red) and p63, showing recombination and no detectable differences in epithelial morphology compared to Cre-negative controls. (C, D) Quantification of epithelial thickness (C) and luminal area (D) in P63-CreER; Stat3$^{fl/fl}$ embryos. (E) Representative transverse sections of Krt5-CreER; Stat3$^{fl/fl}$ (Stat3 KO) embryos stained with H&E and immunofluorescence for TdTomato (red) and KRT5, also showing recombination and comparable epithelial morphology relative to controls. (F, G) Quantification of epithelial thickness (F) and luminal area (G) in Krt5-CreER; Stat3$^{fl/fl}$ embryos. Scale bars: 100µm. Graphs: mean ± SEM; n = 5 embryos per group. Welch's t-test, two-tailed. Three independent experiments were performed per CreER line. The data shown are from one experiment and are representative of all replicates.

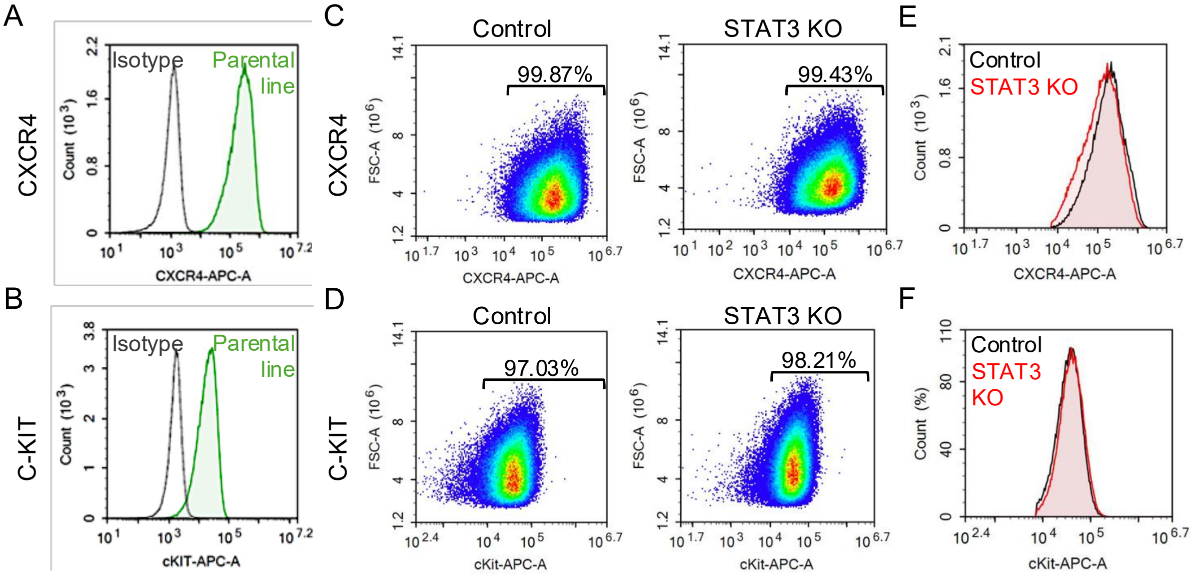


**Supplementary Figure 2. STAT3 is not required for definitive endoderm specification.** (A, B) Flow cytometry analysis of surface markers (A) CXCR4 and (B) c-KIT on day 4 of directed differentiation toward definitive endoderm. Each panel shows overlaid histograms comparing the isotype control (gray shaded) and parental BU3 line (green), demonstrating robust marker expression. (C,D) FSC-A vs. marker expression plots for (C) CXCR4 and (D) c-KIT in control and STAT3 knockout (STAT3 KO) hiPSC lines, showing the percentage of marker-positive cells. (E,F) Overlaid histograms for (E) CXCR4 and (F) c-KIT showing signal intensity (count vs. marker expression) in control (black) and KO (red) cells, demonstrating comparable expression profiles. Three independent experiments were performed, and the data shown are representative of all replicates.

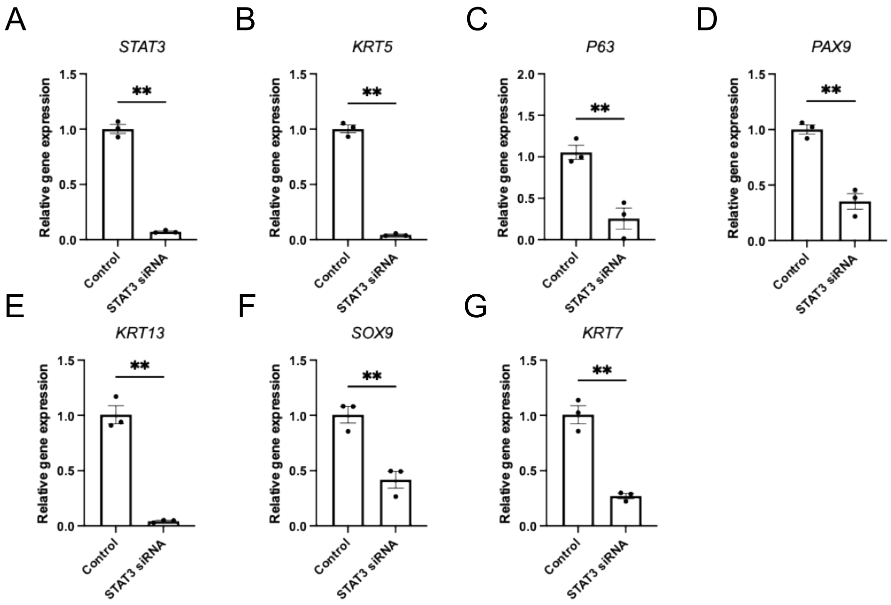


**Supplementary Figure 3. STAT3 regulates epithelial differentiation of hiPSC-derived 3D esophageal organoids.** (A) qPCR analysis confirming STAT3 knockdown in esophageal progenitor cells (EPCs) 24 hours after siRNA transfection (day 24 of differentiation). (B–G) qPCR analysis of epithelial lineage markers in STAT3 siRNA vs. control organoids at 10 days post-embedding, including basal markers (B) *KRT5*, (C) *TP63*, (D) *PAX9*, the suprabasal marker (E) *KRT13*, and fetal esophageal markers (F) *SOX9* and (G) *KRT7.* Graph: mean ± SEM; n = 3 wells per condition. Welch's t-test, two-tailed. **P < 0.01 for all comparisons. Two independent experiments were performed. Data shown are from one experiment and are representative of all replicates.

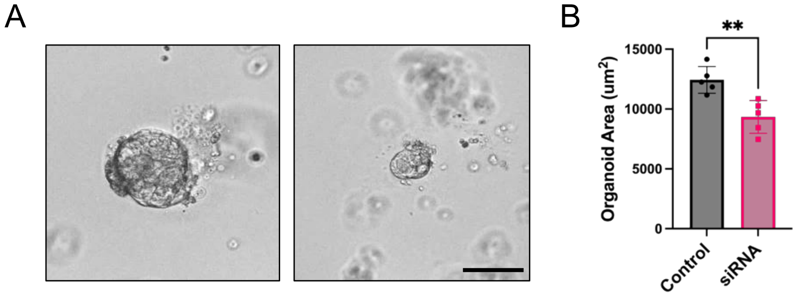


**Supplementary Figure 4. STAT3 regulates growth of primary human fetal esophageal organoids.** (A) Representative brightfield images of primary fetal esophageal organoids derived from 10-week human fetal esophageal epithelium, 7 days after embedding in Matrigel™, transfected with control (scrambled) or STAT3 siRNA. (B) Quantification of organoid size (mean area) across conditions. Scale bar: 200 µm. Graph: mean ± SEM; n = 5 wells per condition, with over 100 organoids measured per well. Welch's t-test, two-tailed. **P < 0.01. One experiment was performed.

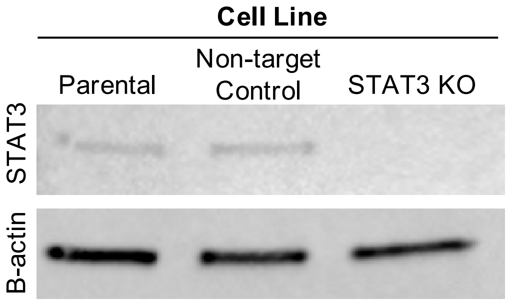


**Supplementary Figure 5. STAT3 knockout in CRISPR-edited hiPSC lines.** (A) Western blot showing loss of STAT3 protein in the STAT3 knockout (STAT3 KO) hiPSC line compared to the scrambled sgRNA (non-target) control and the parental BU3 line. β-actin is shown as a loading control.

A

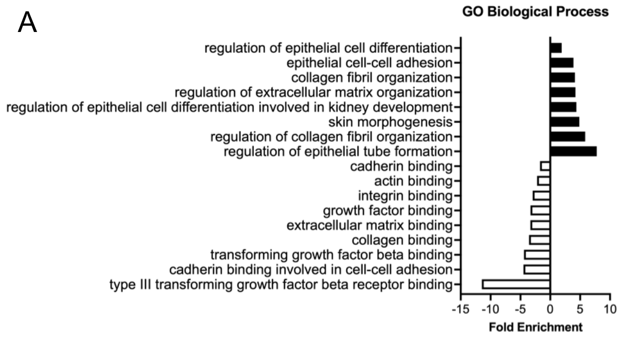


C

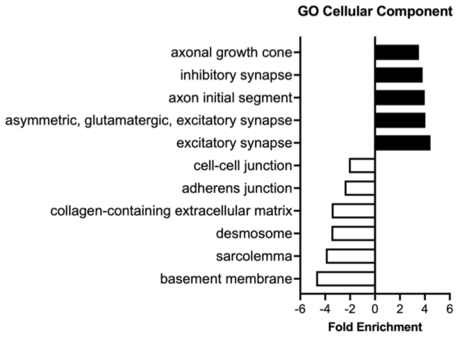


B

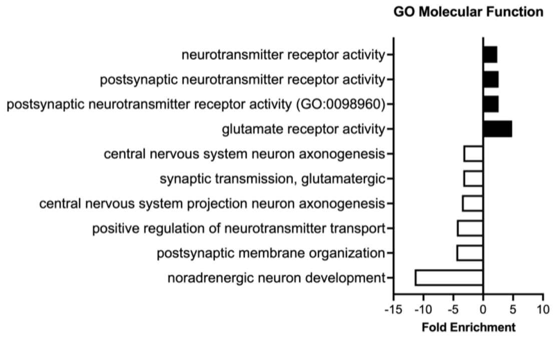


D

| Top 5 upregulated genes in STAT3 KO EPCs |
|---|
| FAR2P1 |
| **FAM107A** |
| **TRIM58** |
| **WNT8B** |
| MED15P9 |

**Supplementary Figure 6. Gene ontology enrichment analysis of differentially expressed genes in STAT3 KO versus control EPCs at day 16 of differentiation.** Enriched (A) biological process, (B) molecular function, and (C) cellular component categories were determined for downregulated and upregulated gene sets (false discovery rate [FDR] ≤ 0.05). Bar plots display representative non-redundant GO terms selected for relevance to epithelial identity, adhesion, and lineage control. (D) Top five upregulated genes in STAT3 KO EPCs ($P_{adj}$ < 0.05 and |log2 fold change| > 1) is included to highlight transcripts of interest for future studies. Gene names in bold have been correlated with developmental signaling and neuronal cytoskeletal remodeling.

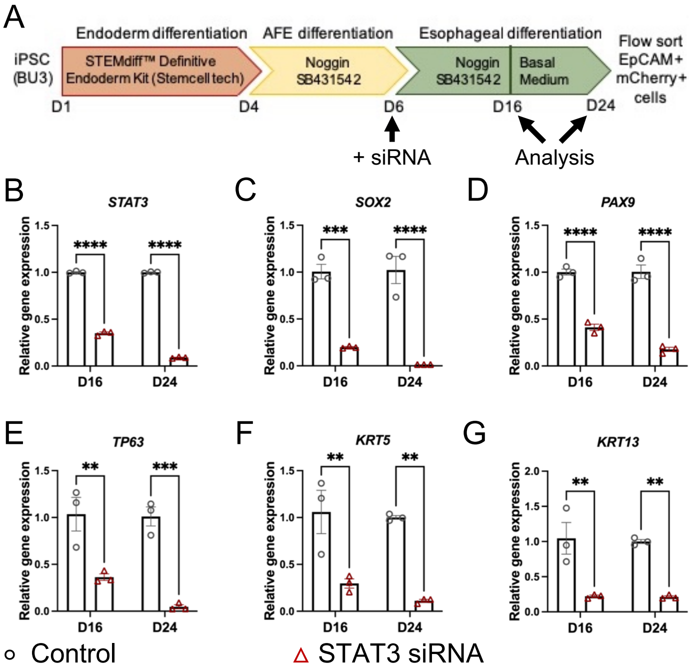


**Supplementary Figure 7. STAT3 knockdown impairs EPC marker expression during directed differentiation.** (A) Schematic of siRNA transfection timeline. AFE cells (day 6) derived from the parental BU3 line were transfected with control or STAT3 siRNA, and gene expression was assessed at days 16 and 24. (B) qPCR analysis confirming sustained suppression of STAT3 expression at both timepoints in STAT3 siRNA-treated cells. (C–G) qPCR analysis of EPC lineage commitment markers (C–E) (SOX2, PAX9, TP63) and (F, G) differentiation markers (KRT5, KRT13) at days 16 and 24. Graph: mean ± SEM; n = 3 wells per condition. Welch's t-test, two-tailed. $^{**}P < 0.01$, $^{***}P < 0.001$, $^{****}P < 0.0001$. Two independent experiments were performed; data shown are from one experiment and are representative of all replicates.

**Supplemental Table 1. List of primary antibodies used for IF, IHC, WB, and FACS**

| Antibody | Calalog Number | Application | Dilution |
|---|---|---|---|
| STAT3 | Cell Signaling Technology, CST9139 | IF, IHC, WB | 1:50 |
| pSTAT3 | Cell Signaling Technology, CST9145 | IHC | 1:100 |
| P63 | Cell Signaling Technology, CST13109 | IF | 1:400 |
| | Cell Signaling Technology, CST39692 | IF | 1:200 |
| SOX2 | Invitrogen, 14-9811-82 | IF, WB | 1:200 |
| Keratin 5 | Biolegend, 905901 | IF | 1:50 |
| | Biolegend, 905503 | IF | 1:50 |
| Ki67 | BD Pharm, 550609 | IF | 1:25 |
| OCT4 | Abcam, ab19857 | IF | 1:200 |
| NANOG | Abcam, ab21624 | IF | 1:1000 |
| FOXA2 | Santa Cruz Biotechnology, sc-374376 | IF | 1:100 |
| SOX17 | R&D Systems, AF1924 | IF | 1:20 |
| Keratin 13 | Abcam, ab92551 | IF | 1:100 |
| TdTomato | MyBioSource, MBS448092 | IF | 1:50 |
| GAPDH | Abcam, ab8245 | WB | 1:2000 |
| beta-Actin | sigma A5316 | WB | 1:5000 |
| EpCAM | Biolegend 324222 | FACS | 5uL per 10 million cells |
| CXCR4 | STEMCELL Technologies, 60089AZ | FACS | 5uL per 10 million cells |
| c-Kit | STEMCELL Technologies, 60087AZ | FACS | 5uL per 10 million cells |

**Supplemental Table 2. List of secondary antibodies used for IF, IHC, and WB**

| Antibody | Calalog Number | Application | Dilution |
|---|---|---|---|
| Donkey anti-Rat | Thermo Fisher Scientific, A48270 | IF | 1:1000 |
| Donkey anti-Mouse | Thermo Fisher Scientific, A32766 | IF | 1:1000 |
| Donkey anti-Rabbit | Thermo Fisher Scientific, A31573 | IF | 1:1000 |
| Donkey anti-Goat | Thermo Fisher Scientific, A21432 | IF | 1:1000 |
| Goat anti-Chicken | Thermo Fisher Scientific, A11039 | IF | 1:1000 |
| Horse anti-Mouse | Vector, BA-2001 | IHC | 1:200 |
| Goat anti-Mouse | LI-COR, 926-68070 | WB | 1:1000 |

**Supplemental Table 3. qPCR primer sequences**

| Gene | Forward Primer | Reverse Primer |
|---|---|---|
| STAT3 | GGTACATCATGGGCTTTATC | TTTGCTGCTTTCACTGAATC |
| TP63 | TTCGGACAGTACAAAGAACGG | GCATTTCATAAGTCTCACGGC |
| SOX2 | CACACTGCCCCTCTCAC | TCCATGCTGTTTCTTACTCTCC |
| PAX9 | GGTGAACGGGTTGGAGAAG | CTGTAGGTCATGTAAGGCGAC |
| IVL | CTGCCTCAGCCTTACTGTG | GCTCCTGATGGGTATTGACTG |
| KRT5 | AGAGCTGAGAAACATGCAGG | AGCTCCACCTTGTTCATGTAG |
| KRT13 | AAGACCATTGAAGAGCTCCG | TGGCATTGTCAATCTCCAGG |
| KRT14 | GAAGTGAAGATCCGTGACTGG | GCAGAAGGACATTGGCATTG |
| YWHAZ | ACTTTTGGTACATTGTGGCTTCAA | CCGCCAGGACAAACCAGTAT |
